# Discrete and conserved inflammatory signatures drive thrombosis in different organs after *Salmonella* infection

**DOI:** 10.1101/2024.01.16.575813

**Authors:** Marisol Perez-Toledo, Nonantzin Beristain-Covarrubias, Jamie Pillaye, Ruby R Persaud, Edith Marcial-Juarez, Sian E. Jossi, Jessica R. Hitchcock, Areej Alshayea, William M. Channell, Rachel E Lamerton, Dean P Kavanagh, Agostina Carestia, William G Horsnell, Ian R. Henderson, Nigel Mackman, Andrew R Clark, Craig N Jenne, Julie Rayes, Steve P. Watson, Adam F. Cunningham

**Author notes:** **Correspondence:** Adam F. Cunningham, B15 2TT, Birmingham, United Kingdom, Steve P. Watson, B15 2TT, Birmingham, United Kingdom. These authors contributed equally to this work (Marisol Perez-Toledo, Nonantzin Beristain-Covarrubias).

## Abstract

Inflammation-induced thrombosis is a common consequence of bacterial and viral infections, such as those caused by *Salmonella* Typhimurium (STm) and SARS-CoV-2. The identification of multi-organ thrombosis and the chronological differences in its induction and resolution raises significant challenges for successfully targeting multi-organ infection-associated thrombosis. Here, we identified specific pathways and effector cells driving thrombosis in the spleen and liver following STm infection. Thrombosis in the spleen is independent of IFN-γ or the platelet C-type lectin-like receptor CLEC-2, while both molecules were previously identified as key drivers of thrombosis in the liver. Furthermore, we identified platelets, monocytes, and neutrophils as core constituents of thrombi in both organs. Depleting neutrophils or monocytic cells independently abrogated thrombus formation. Nevertheless, blocking TNFα, which is expressed by both myeloid cell types, diminished both thrombosis and inflammation which correlates with reduced endothelial expression of E-selectin and leukocyte infiltration. Moreover, tissue factor and P-selectin glycoprotein ligand 1 inhibition impair thrombosis in both spleen and liver, identifying multiple common checkpoints to target multi-organ thrombosis. Therefore, organ-specific, and broad mechanisms driving thrombosis potentially allow tailored treatments based on the clinical need and to define the most adequate strategy to target both thrombosis and inflammation associated with systemic infections.

## Introduction

A key function of inflammation is to provide a rapid response to infection and restrict the dissemination of the pathogen and its components. However, in severe infections, excessive inflammation can also result in the development of a pro-coagulant state that can drive coagulopathies, with deleterious consequences for the host (1). For example, in patients with severe COVID-19, thrombi can form in the lungs, kidneys, and hearts (2, 3) and their presence is associated with poorer patient outcomes in the short- and long-term (4, 5).

Similarly, in humans and animal models of infection, thrombosis and coagulopathies are detected after localized and severe, disseminated bacterial infections, such as those caused by the plague bacillus or *Salmonella* (6–8). Furthermore, a recent study showed that patients with newly diagnosed non-typhoidal salmonellosis (NTS) had an increased risk of developing deep vein thrombosis (DVT) and pulmonary embolism (PE) (9). In patients with typhoid fever, the risk of thrombosis is coincident with thrombocytopenia, high D-dimer levels, increased circulating fibrinogen levels, disseminated intravascular coagulation (DIC), and an increase in markers associated with endothelium activation, such as soluble von Willebrand Factor and endocan (10). Importantly, such changes in thrombosis risk and these coagulation markers do not correlate with the detection of *Salmonella* in the bloodstream as patients have a median detectable burden of only 1 bacterium per ml of blood (11). This suggests that processes activated because of the infection are the main drivers of the changes in coagulation parameters. However, the mechanisms that contribute to thrombosis development after infections such as those caused by *Salmonella* are less clear.

We have previously reported that after *Salmonella* Typhimurium (STm) infection in the mouse, thrombi are detected in the liver from seven days after infection and persist for weeks thereafter (12). Nevertheless, thrombi also form in the spleen within the first day of infection (13), but in this organ, thrombosis resolves rapidly over the next day. Thus, one infection can result in thrombosis in multiple organs with predictable kinetics. In both organs, there is an absolute requirement for clodronate-liposome-sensitive cells (monocyte-lineage cells) for thrombosis. This suggested that the same mechanisms may be shared between both these sites, particularly a need for the cytokine IFN-γ and platelet activation through the podoplanin/CLEC-2 axis, which are key players in this process in the liver (12). However, the discrete kinetics of thrombosis in the spleen suggest that there might also be organ-specific factors that influence this process too, or indeed a core mechanism with some factors shared but other organ-specific factors influencing the induction of thrombosis locally.

To assess this, we have performed a side-by-side analysis of which mechanisms are conserved in the spleen and liver at the times when thrombosis is established in each organ. This analysis reveals that there is a common mechanism that is shared between these organs but there is also a degree of redundancy for some factors between these sites. This suggests that local factors modulate the risk of thrombus development, and this may need to be considered in future strategies to target infection-induced thrombosis.

## Results

### STm-induced thrombosis in the spleen does not require CLEC-2 or IFN-γ

Our previous work showed that thrombosis in the spleen and liver follow different kinetics. Thrombosis in the spleen is detected 24 hours post-infection, whereas in the liver it takes up to 7 days to develop (12, 13). Moreover, the mechanism driving thrombosis in the liver required IFN-γ and the CLEC-2/podoplanin axis (12). Therefore, we decided to focus our analysis on day 1 post-infection (p.i.) for the spleen, and day 7 p.i. in the liver. To evaluate whether the same factors were needed for thrombosis in the spleen, we infected WT and IFN-γ -deficient mice with STm and evaluated thrombosis 24 hours later. IFN-γ-deficient mice had reduced thrombosis in the spleen, but thrombi were still detected in 50% of mice (Fig. 1A and 1B). At this early time point, there was only a minor difference in the bacterial burdens (Supp. Fig. 1A). In contrast, in STm-infected *Clec2^fl/fl^PF4^cre^* mice, which lack the expression of CLEC-2 on platelets and megakaryocytes, thrombosis levels were comparable to levels observed in WT controls (Fig. 1C and 1D). In these mice, the absence of CLEC-2 on platelets had no effect on the bacterial control (Supp. Fig. 1B). These results suggest that, the mechanism of thrombus formation in the spleen is independent of IFN-γ and CLEC-2.

**Figure 1.**
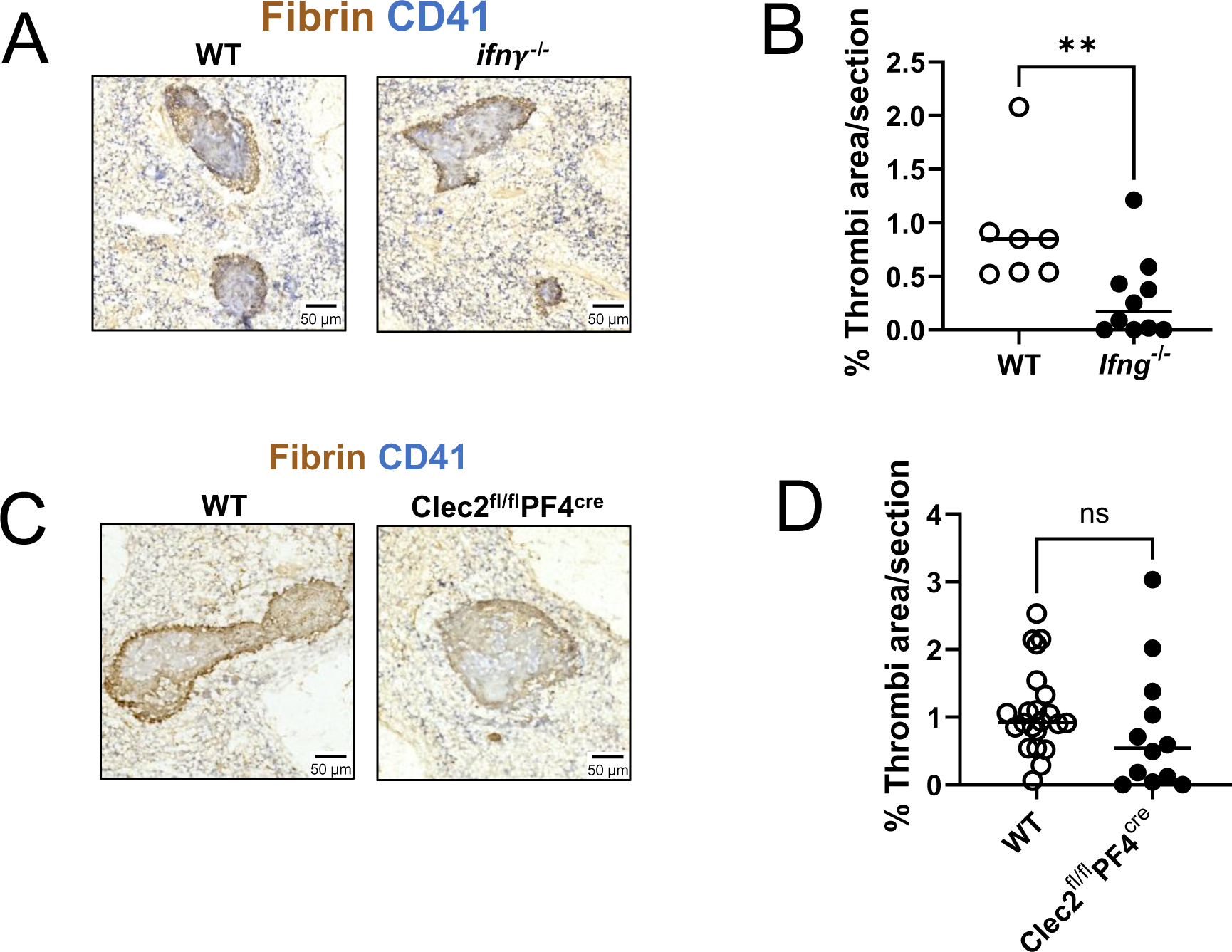
IFN-γ and CLEC-2 are dispensable for thrombosis in the spleen after STm infection. (A) Representative immunohistochemistry of spleen sections from WT and IFN-γ-deficient mice infected with STm for one day. Brown= Fibrin, Blue=CD41. (B) Percentage of area occupied by thrombi in spleens from (A). (C) Representative immunohistochemistry of spleen sections from WT and PF4^Cre^CLEC-2^fl/fl^ mice infected with STm for one day. Brown=Fibrin, Blue=CD41. (D) Percentage of area occupied by thrombi in spleens from (C). Each dot represents an individual mouse. Data was collected from at least two independent experiments. Horizontal lines depict the median. Mann-Whitney with Dunn’s post-hoc test was applied. ** P<0.005, ns=non-significant

### Splenic and liver thrombi contain platelets, monocytes, neutrophils, and fibrin

To better understand the mechanisms of thrombosis in both organs, we evaluated the kinetics of thrombi formation prior to 24 hours post-infection in the spleen, and prior to 7 days in the liver. After infection, thrombi were detected in the spleen by 8 hours and in the liver by 7 days after infection (Fig. 2A). We then analyzed the composition of splenic and liver thrombi by immunohistology. We found that neutrophils (Ly6G) and monocytic cells (Ly6C) were present in both splenic and liver thrombi, alongside with platelets (CD41) and fibrin (Fig. 2B). We then analyzed the change in the frequencies of neutrophils and monocytic cells in spleen and liver after STm infection. In the spleen, the frequency of neutrophils increased from 4 hours post-infection (p.i,), with a drastic increase by 18 hours (Fig. 2C), which coincided with detection of thrombi in this organ (Fig. 2A). In the liver, neutrophils only marginally increased by day 7 p.i., whereas the frequency of monocytic cells increased from day 1 and kept increasing by day 7 (Fig. 2C). To better understand the inter-relationship between these cell populations, we performed intra-vital imaging of the spleen and liver at 24 hours and 7 days after STm infection, respectively. After infection, clusters of monocytic cells, neutrophils and platelets were observed interacting in the splenic vasculature at 24 hours post-infection (Fig. 2D and supp. videos 1-5). In the livers of 7-day infected mice, more neutrophils and monocytic F4/80+ cells were observed in the liver (Fig. 2E and supp. video 4 and 5). The monocytic F4/80+ cells were typically more rounded and had a reduced branched morphology than those observed in non-infected mice, consistent with infiltration of the liver by monocyte-derived macrophages. These results suggest that both neutrophils and monocytic cells respond to STm infection and their presence in the tissues is associated with thrombosis.

**Figure 2.**
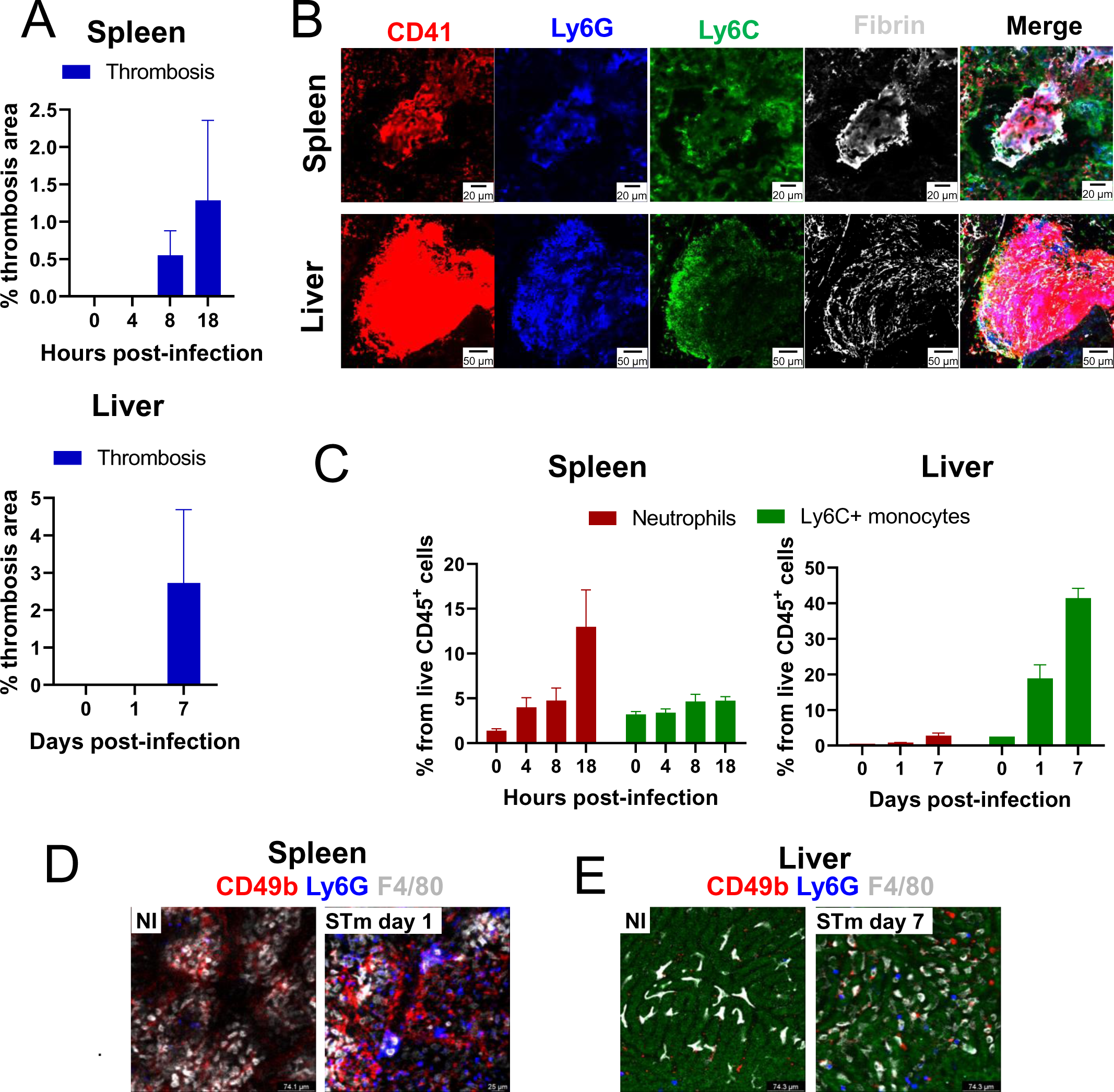
STm infection drives the accumulation of neutrophils and Ly6C^+^ monocytic cells and are found in thrombi. (A) Percentage of the area occupied by thrombi in spleen and liver after STm infection across the indicated time points. (B) Representative images of a spleen thrombus on day 1 (upper row) and a liver thrombus on day 7 (bottom row), stained for CD41 (red), Ly6G (blue), Ly6C (green), and fibrin (grey). (C)Frequency of neutrophils and Ly6C^+^ monocytes from live CD45^+^ cells in spleen (left) and liver (right). Representative fields of view (FOV) of spleen (D) and liver (E) obtained by intra-vital microscopy in non-infected mice (NI) and STm infected mice, 1 day (spleen) or 7 days (liver) post-infection. Cells were stained with CD49b (red), Ly6G (blue) and F4/80 (white). Median ± SEM are depicted.

### Neutrophils are required for thrombosis in the spleen and liver

Since neutrophils are a major component of thrombi in both the spleen and liver (Fig. 2A), we sought to identify whether neutrophils were required for thrombus formation and progression after STm infection. To test this, neutrophils were depleted with a monoclonal anti-Ly6G antibody prior to and during infection with STm (Supp. Fig. 2A), resulting in a reduction in numbers of CD11b^+^Gr1^hi^Ly6C^-^ cells in both the spleen and liver (Fig. 3A). Within the timeframes assessed, neutrophil depletion did not markedly affect bacterial control in either the spleen or liver (Supp. Fig. 3A). Nevertheless, thrombosis was completely abolished in both the spleen and liver in the neutrophil-depleted mice (Fig. 3B and 3C). This suggested that neutrophils are essential for STm-induced thrombosis in both organs. As reported previously (12, 13), mice treated with clodronate liposomes, to reduce monocytic cell numbers, also had negligible levels of thrombosis (Fig. 3D and 3E; Supp. Fig. 2B and 3B). Together, these results show that both neutrophils and monocytic cells are independently required for thrombosis in both spleen and liver.

**Figure 3.**
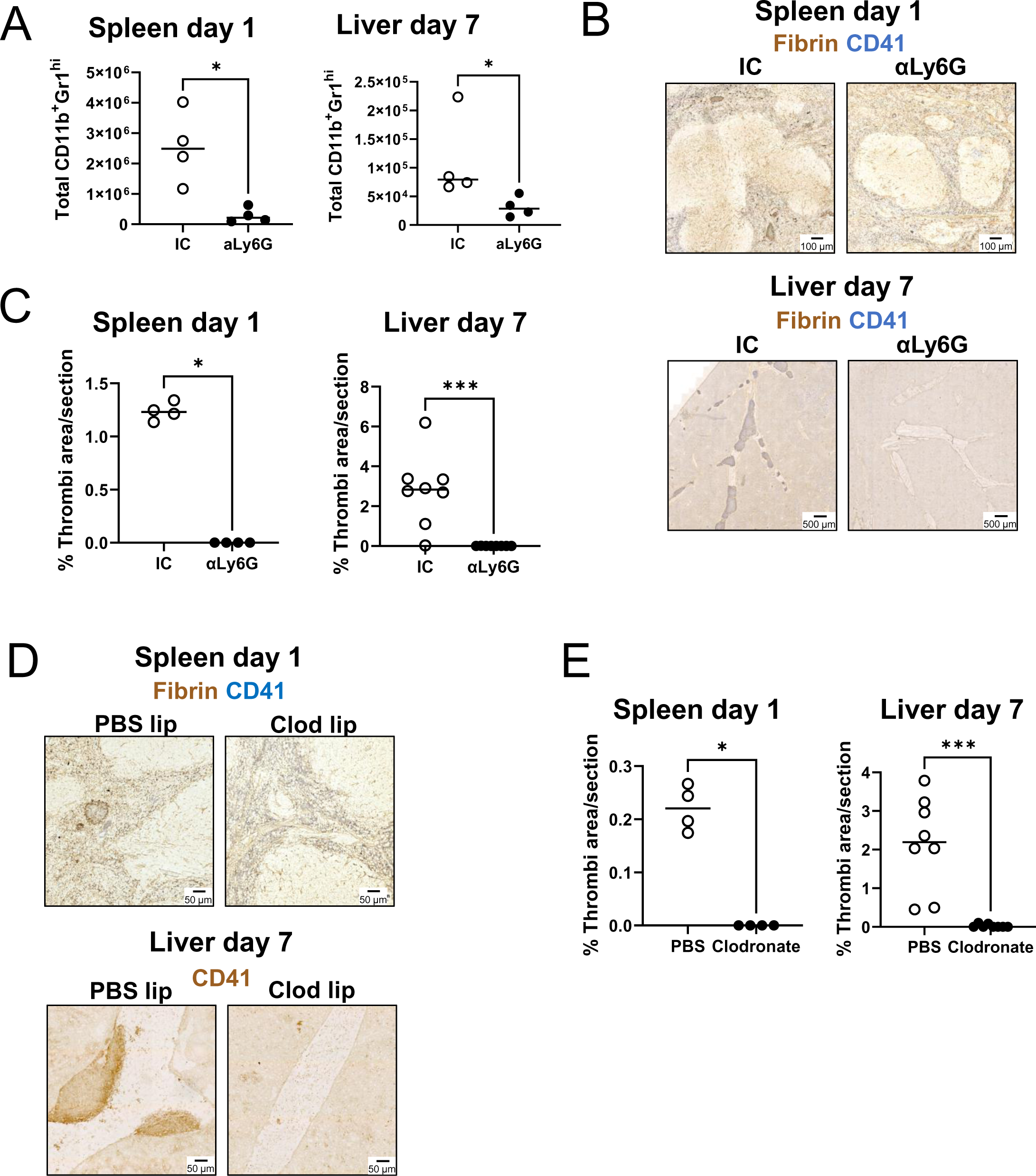
Neutrophils and monocytic cells are needed to drive thrombosis in spleen and liver after STm infection. (A) Total number per organ of CD11b^+^Gr1^hi^ cells in isotype control treated (IC) or anti-Ly6G treated mice in the spleen (1 day post-infection) or liver (day 7 post-infection). (B) Representative immunohistochemistry (Fibrin= brown, CD41=blue) of spleen and liver sections from isotype control (IC) or anti-Ly6G treated mice. (C) Quantification of the area occupied by thrombi in spleen and liver from (B). (D) Representative immunohistochemistry (Fibrin= brown, CD41=blue) of spleen and liver sections from PBS liposomes or clodronate liposomes treated mice (E) Quantification of the area occupied by thrombi in spleen and liver from (D). Each dot represents an individual mouse. Representative data from at least two independent experiments. Horizontal lines depict the median. Mann-Whitney with Dunn’s post-hoc test was applied. * P<0.05, ***P<0.001

### TNFα drives thrombosis in the spleen and liver after STm infection

The requirement for monocytic cells and neutrophils suggested that inflammatory cytokines contribute to thrombosis. One major cytokine involved in the early response to STm is TNFα, which can activate endothelial cells to influence leukocyte migration. After infection, intracellular TNFα was readily detected in both neutrophils and Ly6C^+^ monocytic cells in the spleen and liver after 1 day and 7 days post-infection, respectively (Fig. 4A). At day 1 post-infection, most staining for TNFα was associated with F4/80^+^ cells in the spleen (Fig. 4B).

**Figure 4.**
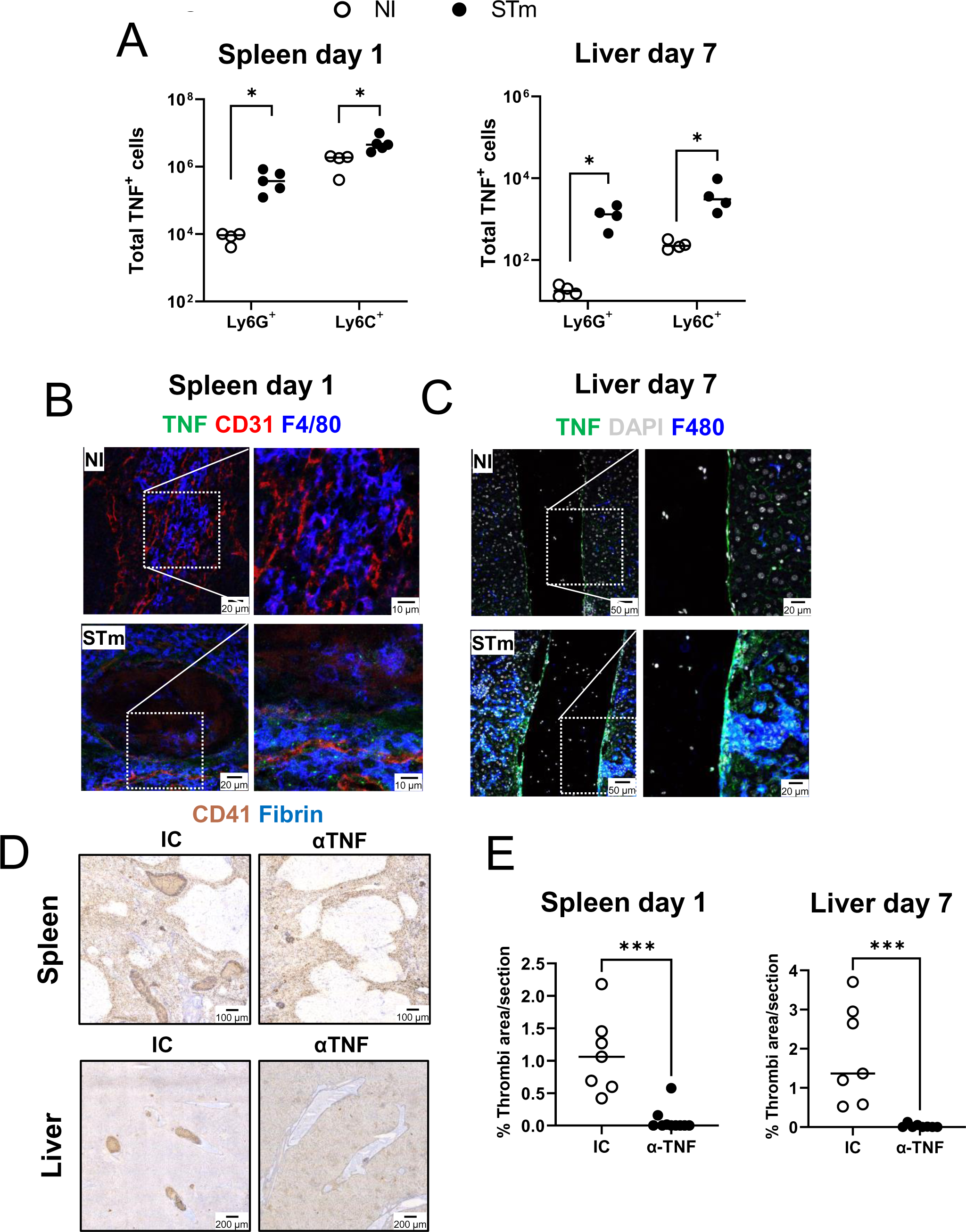
Blocking TNFα prevents thrombosis after STm infection. (A) Total number per organ of TNFα producing neutrophils (CD11b+Ly6G+) and Ly6C+ monocytes (CD11b+ Ly6C+) in the spleen (1 day post-infection) and liver (7 days post-infection). (B) Representative images of spleen sections stained for TNFα (green), CD31 (red) and F4/80 (blue). (C) Representative images of liver sections stained for TNFα (green), DAPI (grey) and F4/80 (blue). Images to the right correspond to magnifications from the areas delimited by the white dotted lines. (D) Experimental approach to neutralize TNFα during STm infection. (E) Representative images from sections stained by immunohistochemistry (Fibrin= blue, CD41=brown) (F) Quantification of the area occupied by thrombi in spleen and liver from (E). Each dot represents an individual mouse. Representative data from at least two independent experiments. Horizontal lines depict the median. Mann-Whitney with Dunn’s post-hoc test was applied.

Moreover, clear TNFα staining was observed around the blood vessels of livers 7 days after infection (Fig. 4C). To test whether TNFα neutralization could prevent thrombosis developing after STm infection, mice were treated with an anti-TNFα antibody and the levels of thrombosis evaluated on days 1 and 7 post-infection (Supp. Fig. 2C). Blocking TNFα reduced the development of thrombosis in the spleen and the liver to near undetectable levels (Fig. 4D, E). The administration of anti-TNFα antibodies had no impact on bacterial numbers on day 1 post-infection, and only a modest impact at day 7 (Supp. Fig. 3C).

One of the effects of TNFα is the upregulation of adhesion molecules on the endothelium, such as CD62E (14). In non-infected mice, CD62E was not detectable in the vasculature in either the spleen or the liver (Fig 5A and 5B). However, after infection, CD62E was upregulated in the spleen and liver at days 1 and 7, respectively. In mice receiving anti-TNFα antibodies, less CD62E expression was detected in the vasculature (Fig. 5A-D). To further confirm that the increase of CD62E expression is through TNFα signaling, spleen and liver sections from mice deficient in TNFα receptors (*p55^-/-^/p75^-/-^*) were stained for CD62E (Supp. Fig. 4A). In contrast to WT mice, TNF receptor-deficient mice did not have detectable CD62E staining after infection, nor did they have any detectable thrombi. One consequence of these effects of TNFα on the endothelium could be a reduction in the numbers of immune cells in the tissues. Flow cytometry showed that TNFα treatment resulted in a selective reduction in the numbers of neutrophils in the spleen compared to isotype control-treated mice (Fig. 5E), although Ly6C+ monocytic cells increased modestly. In the liver, TNFα blocking resulted in reduced numbers of Ly6C+ monocytic cells compared to isotype control treated mice, but no effect was seen in the numbers of neutrophils (Fig. 5F). These results suggest that TNFα promotes thrombosis by inducing upregulation of CD62E and leukocyte migration.

**Figure 5.**
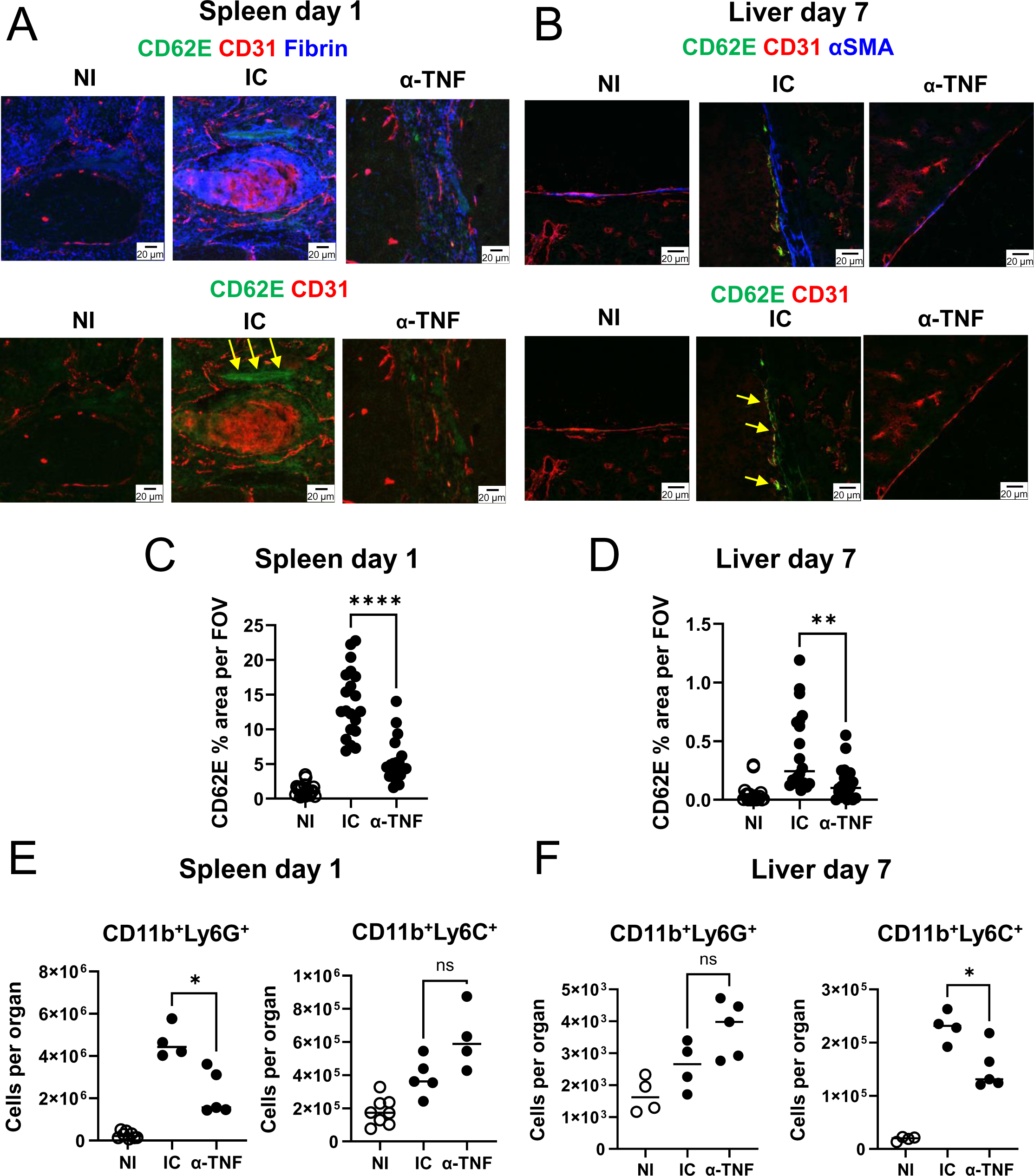
TNFα induced by STm infection drives CD62E expression. (A) Representative immunofluorescence images from spleen 1 day post-infection stained for CD62E (green), CD31 (red), fibrin (blue). (B) Representative immunofluorescence images from liver 7 days post-infection stained for CD62E (green), CD31 (red), α-SMA (blue). NI= non-immunized, IC= Isotype control, α-TNF= anti-TNFα. Yellow arrows indicate areas of positive CD62E staining. Frequency of CD62E positive area per field of view (FOV) in spleen (C) and liver (D) from (A) and (B). Each dot represents a different FOV. Frequency of CD11b^+^Ly6G^+^ cells (neutrophils) and CD11b^+^Ly6C^+^ cells (Ly6C^+^ monocytes) in spleen (E) and liver (F) in non-immunized (NI), isotype control (IC) or anti-TNFα treated mice. Each dot represents an individual mouse. Representative data from at least two independent experiments. Horizontal lines depict the median. Mann-Whitney with Dunn’s post-hoc test was applied. * P<0.05, **P<0.005, ***P<0.001.

### Tissue factor is upregulated after STm infection and is essential for thrombosis

An additional effect of TNFα on the vasculature is the upregulation of tissue factor (TF) on endothelial cells (15, 16). Moreover, cells like neutrophils and monocytes can contribute as sources of TF (17). Because fibrin is a major component of splenic and liver thrombi (Fig. 2B), and fibrin deposition results from the activation of thrombin that can be initiated by TF, we investigated the role of TF in the development of STm-induced thrombosis. Imaging showed that TF was increased in the spleen and liver, 1 day and 7 days post-infection, respectively (Fig. 6A). TF was detected in the periphery of thrombi, with a similar staining pattern to fibrin (Fig. 6B) and co-stained with Ly6G+ neutrophils (Fig. 6C). TF was also detected in perivascular cells, but staining did not associate with CD31+ cells (Fig. 6D). TF promotes the generation of fibrin through thrombin activation. To determine the presence of active thrombin after STm infection, we performed intra-vital microscopy and incorporated a probe that emits fluorescence when cleaved by active thrombin (18). Elevated thrombin activity was detected in the splenic vasculature on day 1 after STm when compared to non-infected mice (Fig. 6E and 6F). Similar experiments in the liver on day 7 post-infection showed thrombin activity was increased in the vasculature of STm-infected mice compared to non-infected mice (Fig. 6E and 6F). Furthermore, active thrombin was detectable in thrombi (Fig. 6G). Finally, the requirement of TF for thrombosis after STm infection was assessed by infecting mice that express low levels of TF (mTF^-/-^, hTF^+/+^) alongside heterozygous control mice (mTF^+/-^, hTF^+/-^) (19). Thrombi were not detected in either the spleen or the liver of mTF^-/-^, hTF^+/+^ mice (Fig. 5H and 5I), despite both groups having similar bacterial burdens (Supp. Fig. 3D). These results suggest that, upon STm infection, TF is required for thrombosis post-STm infection.

**Figure 6.**
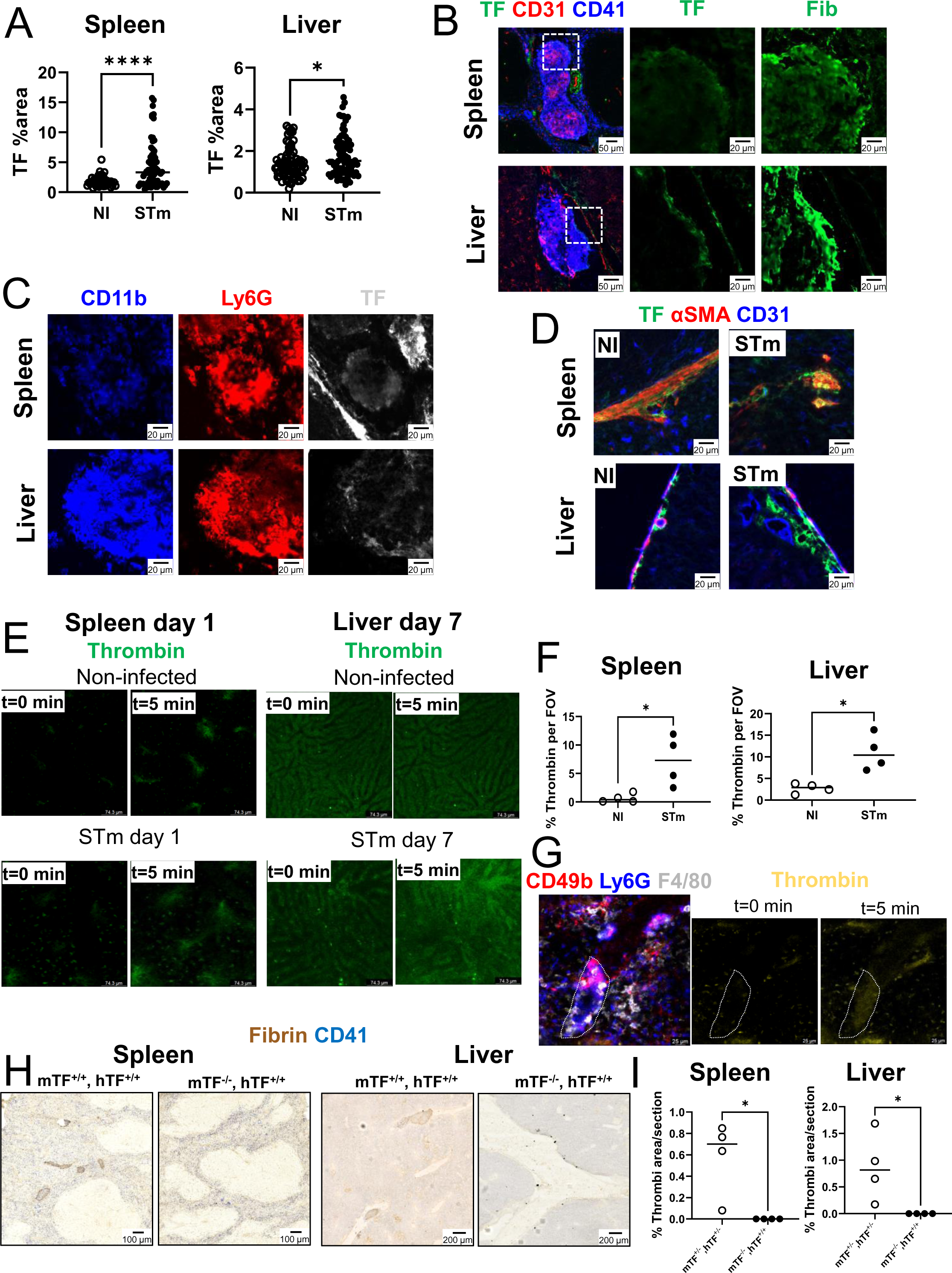
STm induced thrombosis is driven by tissue factor. (A) Quantification of tissue factor positive pixel area in spleen and liver sections from non-infected controls (NI) or STm infected mice, 1 day post-infection (spleen) and 7 days post-infection (liver). Each dot represents a different field of view. (B) Representative images of spleen and liver thrombi stained for tissue factor (TF, green) or fibrin (Fib, fibrin), CD31 (red) and CD41 (blue). (C) Representative images of spleen and liver thrombi stained for tissue factor (TF, white), Ly6G (red) and CD11b (blue). (D) Representative images of spleen and liver sections from non-infected controls (NI) or STm infected mice stained for tissue factor (TF, green), α-SMA (red), and CD31 (blue). (E) Representative fields of view (FOV) of thrombin activation in spleen and liver obtained by intra-vital microscopy. T=0 min shows the first frame of the recording before the administration of the thrombin probe. T=5 min shows the frame in the same FOV 5 minutes after administration of the thrombin probe. (F) Quantification of thrombin-positive pixels after 5 min of administration of the thrombin probe. (G). Representative image of a splenic thrombus captured by intra-vital microscopy, on t=0 min and t=5 min after the administration of the thrombin probe (yellow). Dashed lines delimitate the thrombus. (H) Representative images of spleen (left) and liver (right) sections stained for fibrin (brown) and CD41 (blue) in heterozygous mice (expressing human TF and mouse TF, mTF^+/-^, hTF^+/-^) and mice with low tissue expression (mTF^-/-^, hTF^+/+^). (I) Quantification of the area occupied by thrombi in spleen and liver from (G). Each dot represents an individual mouse. Horizontal lines depict the median. Mann-Whitney with Dunn’s post-hoc test was applied. * P<0.05, ***P<0.001, P<0.0001

### Inhibition of PSGL-1 prevents thrombosis by limiting neutrophil and monocyte recruitment

TF can accumulate in thrombi via a mechanism that involves P-selectin Glycoprotein Ligand-1 (PSGL-1) (20). Dual staining for TF and PSGL-1 identified co-localization of these factors within thrombi (Fig. 7A). In models of DVT, peptides that block PSGL-1 function can prevent thrombosis (21). Therefore, we tested whether blocking PSGL-1 prevented thrombosis after STm infection. We administered an anti-PSGL-1 antibody and evaluated thrombi in the spleen and liver at day 1 and day 7 post-infection, respectively (Supp. Fig. 2D). At these timepoints, fewer thrombi were detected in the spleens and livers of mice receiving anti-PSGL1 (Fig. 7B and 7C). Blocking PSGL-1 did not affect bacterial control at either of the time points tested (Supp. Fig. 3E). Microscopy and flow cytometry showed that after PSGL-1 blockade, fewer neutrophils were detected in the spleen 24 h after infection, whilst the frequency of Ly6C^+^ monocytic cells remained unaffected (Fig. 7D and 7E). In contrast, in the liver at day 7 post-infection fewer CD11b^+^ Ly6G^-^ cells were detected, but frequencies and numbers of CD11b^+^Ly6C^+^ cells were similar between the two groups (Fig. 7F and 7G). These results show that PSGL-1 has a role in promoting thrombosis in both the spleen and liver.

**Figure 7.**
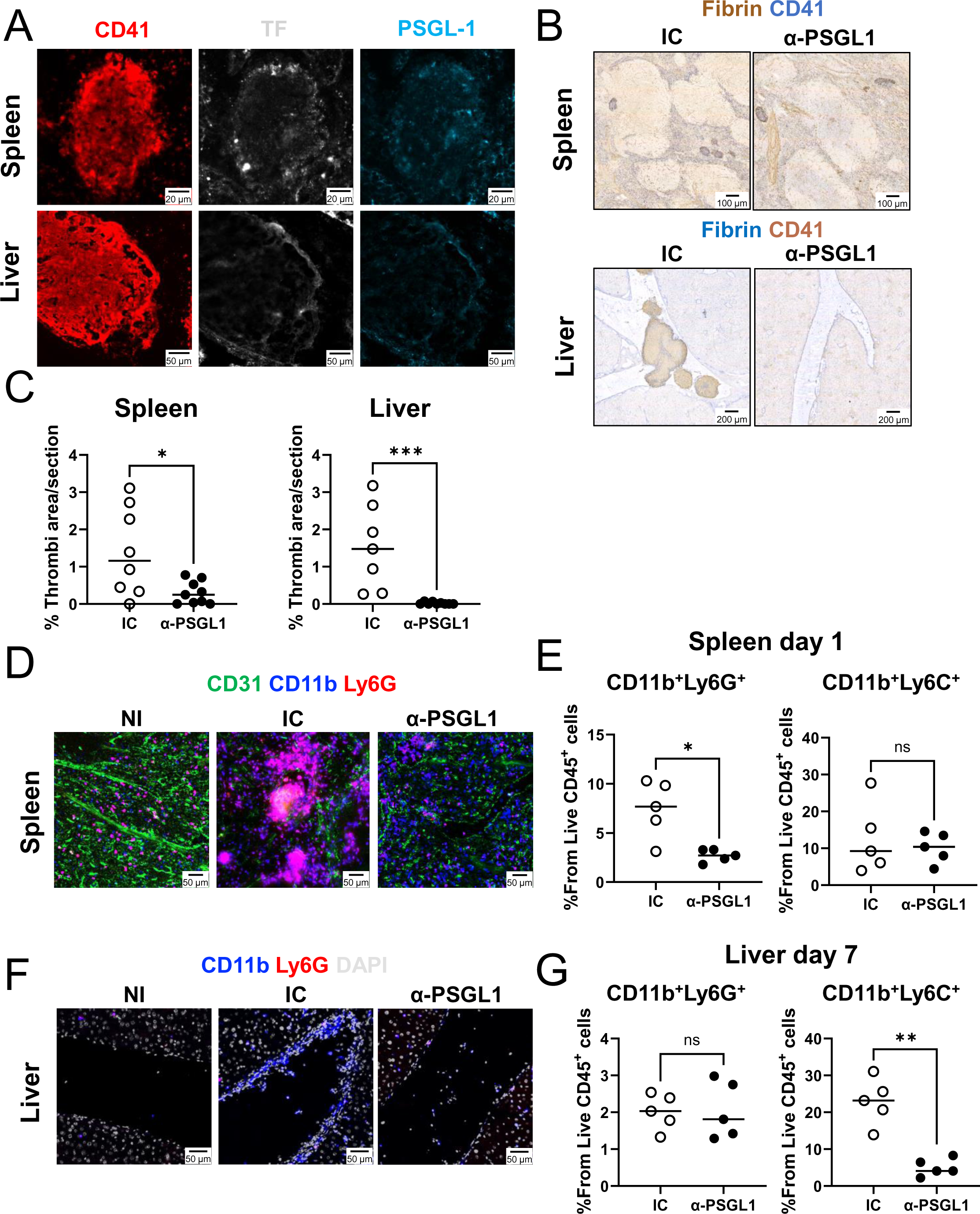
PSGL-1 prevents thrombosis by modulating neutrophil and monocyte recruitment. (A) Representative images of spleen (top row) and liver (bottom row) sections from mice infected with STm at day 1 p.i. (spleen) and day 7 p.i. (liver), stained for CD41 (red), tissue factor (TF, white), and PSGL-1 (blue). (B) Representative images of spleen (top row) and liver (bottom row) sections from mice infected with STm and treated either with isotype control (IC) and anti-PSGL1 (α-PSGL1), stained for fibrin (blue) and CD41 (blue). (C) Quantification of the area occupied by thrombi in spleen at day 1 p.i., or liver at day 7 p.i., from mice treated with isotype control (IC) or anti-PSGL1(α-PSGL1). (D) Representative images of spleen sections stained for CD31 (green), CD11b (blue), and Ly6G (red) (E) Frequency of neutrophils (CD11b^+^Ly6G^+^) or Ly6C^+^ monocytes (CD11b^+^Ly6C^+^) quantified by FACSs in spleen of isotype control treated (IC) or anti-PSGL1 treated mice (α-PSGL1) 1 day p.i. (F) Representative images of liver sections stained for CD11b (blue), Ly6G (red) and DAPI (grey). Sections were obtained from non-infected mice (NI), isotype control treated mice (IC) or anti-PSGL-1 treated mice at day 7 p.i. (liver). (G) Frequency of neutrophils (CD11b^+^Ly6G^+^) or Ly6C^+^ monocytes (CD11b^+^Ly6C^+^) quantified by FACSs in livers of isotype control treated (IC) or anti-PSGL1 treated mice (α-PSGL1) 7 days p.i. Each dot represents a single mouse. Data was collected from at least two independent experiments. Mann-Whitney with Dunn’s post-hoc test was applied. * P<0.05, **P<0.005, ***P<0.001.

## Discussion

A potential consequence of serious infections is the development of thrombosis. The COVID-19 pandemic demonstrated the clinical importance of understanding the mechanisms and effects of infection triggered thrombosis. In this work, we have shown that thrombosis in the mouse model of *Salmonella* infection contains a common pathway that also has organ-specific modulating factors. The common pathway between the organs requires neutrophils, monocytic cells, TNFα and TF activity, whereas the most prominent difference between these organs is the differential requirement for IFN-γ and CLEC-2. This suggests discreet thrombogenic pathways that are organ-specific, but that converge in a common mechanism involving inflammation and coagulation.

*Salmonella*-induced splenic and liver thrombi contain platelets, neutrophils, monocytes, and fibrin. The importance of neutrophils and monocytes is demonstrated by the finding that both cell types are needed to support thrombus formation. Additionally, intravital microscopy identified the close interactions between both leukocyte subsets, platelets, and the blood vessels. In the spleen, cellular aggregates consistent with thrombi morphology were observed and had a similar presentation as thrombi imaged using histology. Although technical limitations restricted our study of the portal vein in the liver, where thrombi are mostly found, interactions between neutrophils and monocytic cells in the sinusoids were observed and indicates that similar interactions can occur in bigger blood vessels. Moreover, there is a co-requirement for neutrophils and monocytes for thrombosis development in both spleen and liver, although there may be subtle differences in their roles depending on the organ. For example, our results indicate that neutrophils play a major role for thrombosis in the spleen, whereas in the liver, monocytes may play a more dominant role. This possibility is supported by the observation that blocking TNFα and PSGL-1 reduced thrombosis in both organs but had distinct effects on the numbers of neutrophils and monocytes in either organ. Nevertheless, a recent report showed that clodronate administration also impairs neutrophil function, without affecting cell viability (22). Hence, we cannot rule out that the results of our study with clodronate are not due in part to a collateral effect on neutrophils.

The differing requirements for IFN-γ and CLEC-2 between the liver and spleen was unexpected. Using a model of inferior vena cava ligation, others have shown that mice with a deficiency in CLEC-2 are protected from developing DVT (23). Our results suggest that not all mechanisms of thrombosis are conserved at all vascular sites throughout the host.

Therefore, the mechanisms that are identified in one site may not be generalizable to another vascular site where thrombi form. Other factors could contribute to the redundancy for IFN-γ and CLEC-2 that we have observed for the spleen. For instance, it may relate to the different timings of thrombosis in the different organs. Alternatively, it may reflect the higher density of monocytic-lineage cells and neutrophils present in the spleen at the time of pathogen encounter and as such there may be a lower threshold for inducing thrombosis in this organ, thus making the podoplanin-CLEC-2 pathway redundant.

Monocytes and neutrophils could promote thrombosis after STm through the production of TNFα, which is necessary for thrombosis and is produced by both cell types after STm infection (24). Nevertheless, blocking TNFα had a modest impact on bacterial control, which is consistent with the role of TNFα in granuloma formation, function and tissue organization (25). TNFα-blocking agents are widely used to treat multiple inflammatory diseases, including psoriasis, Crohn’s disease, and rheumatoid arthritis (Reviewed in (26)). Still, anti-TNFα therapy in sepsis has only shown a modest increase in the survival (27). However, an increased risk of infection, such as those caused by *M. tuberculosis*, has been reported previously in patients under anti-TNFα treatment (28). Therefore, an agent that can block the pathological role of inflammation, without impacting the effector branch for bacterial control, is needed in pathologies derived from infection-triggered inflammation. Thus, targeting molecules such as PSGL-1 may be promising in such situations. Indeed, blocking PSGL-1 molecules can moderate organ damage in models of sepsis (29–31). Nevertheless, it should be noted that although PSGL-1-deficient mice orally infected with STm did not have increased bacterial burdens in the spleen, they did have more severe infections in the colon and overall, a modest reduction in survival, suggesting this strategy may not apply to all types of infection (32).

Infection is an essential step for inducing thrombosis, yet thrombi themselves contain a paucity of bacteria (13). Similarly, bacteria are present in both the spleen and liver at similar and near-peak levels in both organs throughout the first week of infection. Furthermore, both neutrophils and monocytic-lineage cells are essential for thrombosis to develop. Despite these parallel features, the distinct kinetics suggest that other factors must contribute to drive thrombosis in specific sites with these kinetics. One of the contributing factors could be the local activation of the vasculature, which could provide a critical component that “zip codes” where a thrombus will form. Therefore, we propose a model where the priming factor is bacterial infiltration into tissues, in which the bacterium resides in cells or niches proximal to the blood vessels. The resulting local secretion of TNFα results in the activation of the vasculature and the upregulation of CD62E, which promotes the recruitment of neutrophils or monocytic-lineage cells through the blood, and the interaction of these cells with the vasculature, adjacent to where the bacteria are localized. In the case of the liver, activation of the podoplanin/CLEC-2 axis would then contribute to local platelet activation. Alongside, exposure and activation of TF would then promote fibrin deposition and eventual thrombus formation (Outlined in the model shown in Fig. 8).

**Figure 8.**
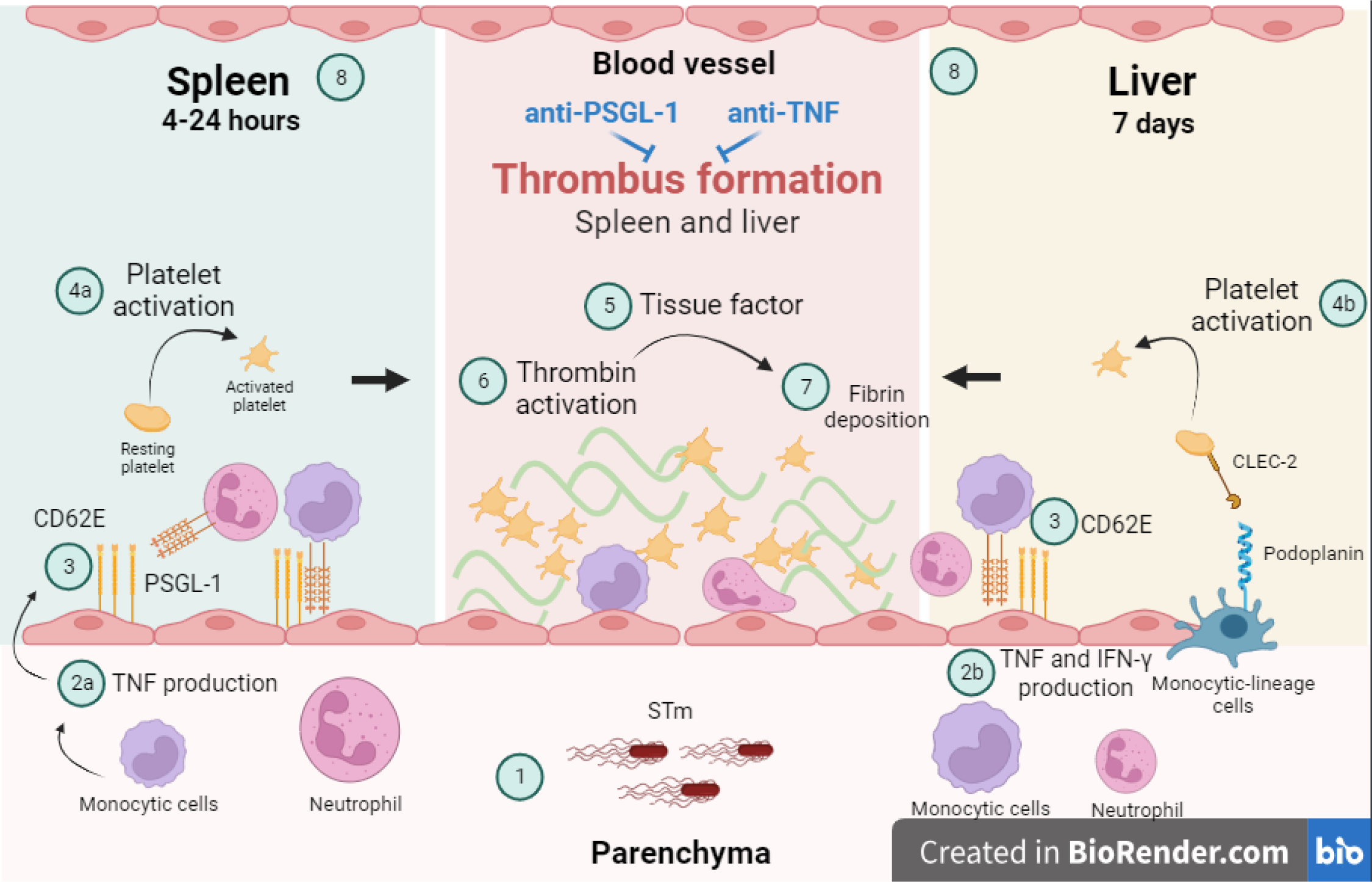
Proposed model for induction of STm-induced thrombosis in the spleen and liver. **Spleen:** STm invades the tissues including peri-vascular sites (1). This results in the local production of TNFα (2) and the further recruitment of phagocytes to parenchymal sites and proximal vessels. PSGL-1 and the upregulation of CD62E on the vasculature mediate neutrophil recruitment to the spleen and phagocyte interactions in the vasculature (3). Activation of platelets in the spleen occurs through an unknown mechanism but may result from the high concentration of intra-organ innate phagocytes (4a). The combination of TF expression of the subendothelium (5) and the activation of thrombin (6), which promotes fibrin deposition (7) results in thrombus development (8). **Liver:** The pathway to thrombosis in the liver is largely conserved except for the additional critical need for IFN-γ (2b), which enhances local monocytic-lineage cell recruitment and the requirement for platelet expression of CLEC-2 (4b) which is required for platelet activation and thrombus development. Finally, this process can be stopped in both organs by the administration of anti-PSGL-1 antibodies or anti-TNFα (8).

## Methods

### Study approval

Mice were used in accordance with the Home Office guidelines at the Biomedical Services Unit of the University of Birmingham under the project License P0677946.

### Mice and infection protocol

C57Bl/6J mice from 6-8 weeks of age were purchased from Charles River. Low Tissue Factor mice, IFN-γ-deficient mice (33), PF4^Cre^CLEC-2^fl/fl^ mice (34), and TNF receptor deficient mice (*p55^-/-^p75^-/-^*) (35) were bred and maintained at the Biomedical Service Unit of the University of Birmingham. Low Tissue Factor mice were phenotyped as described elsewhere (19).

Infection was performed by injecting i.p. 5×10^5^ CFU of an attenuated *Salmonella* Typhimurium strain *ΔaroA* SL3261. In some experiments, monoclonal antibodies were administered during or before the infection to block or deplete specific cell types. Briefly, 500 μg per mouse of an anti-Ly6G antibody (Clone 1A8, Bioxcell), or 300μg of a blocking anti-TNFα antibody (Clone XT3.11, Bioxcell) were injected i.p. prior to infection, and on day 3 and 5 post-infection (Supp Fig. 2A and C). The same amount of rat IgG (Sigma) was used as isotype control. To deplete monocytic cells, 200 µL of clodronate liposomes or PBS liposomes were injected i.p. before infection, and on day 3 and 5 post-infection as described before (Supp Fig. 2B) (12, 13). To block PSGL-1, 200 μg of the clone 4RA10 (Bioxcell) was injected prior infection for the studies in the spleen, and on day 3 and 5 post-infection for studies in the liver (Supp Fig. 2D).

### Intra-vital Microscopy

Male C57Bl/6 mice were infected as described above. 24 hours or 7 days after infection, mice were anesthetized with a mixture of ketamine hydrochloride (200 μg/kg) and xylazine (10mg/kg) and liver and spleen imaged as reported elsewhere (36). Anti-mouse CD49b (1.6 µg, clone HMα2, BioLegend), anti-mouse F4/80 (1.6 µg, clone BM8, BioLegend), anti-Ly6G (1.6 µg, clone 1A8, BioLegend) were injected i.v. 30 min prior imaging. To visualize active thrombin, mice were prepped as described above, and after imaging was started, a fluorescent probe was injected i.v. (Anaspec). 5 to 10-minute-long videos were recorded using an inverted Leica SP8 microscope (Leica Microsystems). Image analysis was performed with Fiji.

### Immunohistology

5 µm cryosections of spleens and livers were fixed in acetone for 20 min and stored at -20 °C until analysis. Immunohistochemistry (IHC) was performed as described previously (13). Briefly, sections were re-hydrated in Tris-Buffered Saline pH 7.6 at room temperature and stained in the same buffer with primary antibodies for 45 minutes. HRP-conjugated or biotin-conjugated secondary antibodies and ABComplex alkaline phosphatase (Dako) were used as secondaries. HRP activity was detected with SIGMA*FAST* 3-3’Diaminobenzidine tablets (Sigma-Aldrich), whereas alkaline-phosphatase activity was detected using naphtol AS-MX phosphate and fast blue salt with levamisole (All from Sigma-Aldrich). For immunofluorescence, sections were re-hydrated in PBS pH 7.4 and blocked for 10 minutes with 10% fetal bovine serum (FBS) in PBS. For the liver, additional biotin blocking steps were performed prior staining with an avidin/biotin blocking kit (Vector Laboratories) following the manufacturer’s instructions. Incubation with antibodies was carried out in the dark at room temperature for 40 minutes. Slides were mounted in Prolong Diamond (ThermoFisher) and curated during 24 hours at room temperature before imaging. A detailed list of antibodies used can be found in supplementary table I

### Flow Cytometry

Single-cell suspensions from spleens were prepared by mashing approximately 20 mg of tissue through a 70 μm cell strainer (Falcon). Red blood cells were lysed with ACK Lysis buffer (Gibco) and cells resuspended in RPMI with 10% FBS. Liver single-cell suspensions were prepared as described previously (12). Cells were stained for viability with Zombie Aqua (Biolegend, UK) prior incubation with primary antibodies during 30 minutes at 4°C. For intracellular staining of TNFα, 5 x 10^6^ cells were seeded in 48-well plates, stimulated with 5 μg/mL of heat-killed STm, and incubated at 37°C with 5% CO_2_, in the presence of Golgi Stop (BD Biosciences). After overnight incubation, extracellular staining was performed as described above. Intracellular staining was performed at room temperature using the BD Cytofix/CytopermTM fixation/permeabilization kit (BD Biosciences) according to the manufacturer’s instructions. Data acquisition was performed with a CytoFLEX using the CytEXPERT software (Beckman Coulter), and data was analyzed with FlowJo Software v10.5 (Tree Star). Neutrophils were defined as CD11b^+^Ly6G^+^, and monocytes were defined as CD11b^+^Ly6G^-^CD3^-^B220^-^NK1.1^-^Ly6C^+^. The proportion of these cells is expressed as frequency from live CD45^+^ leukocytes. A list of all reagents used can be found in supplementary table I.

### Statistical analysis

The two-tailed Mann-Whitney non-parametric sum of ranks test was used to determine statistical significance. The p values were calculated using GraphPad Prism version 10.1.0 and these were interpreted as significant where the p value was ≤ 0.05.

### Data availability

Values for all data points in graphs are reported in the Supporting Data Values file. Additional data are available from the corresponding author (AFC and SPW), upon reasonable request.

## Supporting information

Supp video 1. Spleen STm 24 hours p.i.

Supp video 2. Spleen STm 24 hours p.i.

Supp video 3. Spleen Stm 24 hours p.i.

Supp video 4. Spleen non-immunised

Supp video 5. Spleen STm 6 hours p.i.

Supp video 6. Liver D7

Supp video 7. NI Liver

## Author contributions

MPT and NBC conceived and designed the analysis, performed the experiments, and wrote the paper. JP, RP, EMJ, SEJ, JRH, AA, WMC, RL, AC and DK performed experiments. WGH, IRH, NM, AC, CJ contributed with reagents. JR, SPW and AFC discussed the results and supervised the project. All authors contributed to the final version of the manuscript.

## Acknowledgments

The authors would like to thank the personnel of the Biomedical Service Unit, the University of Birmingham Flow Cytometry Services, and the Microscopy Facilities from the University of Birmingham for their support. The authors would also like to thank Ms. Laura Godin and Dr. Fien von Meijenfeldt for their assistance with experiments. This work was funded by a grant by the Medical Research Council (MR/N023706/1) awarded to AFC. REL was supported by a scholarship awarded by the Wellcome Trust. MPT was supported by a Global Mobility Award by the Wellcome Trust. SPW is a British Heart Foundation chair (CH/03/003).

## References

1. Beristain-Covarrubias N, Perez-Toledo M, Thomas MR, Henderson IR, Watson SP, and Cunningham AF. Understanding Infection-Induced Thrombosis: Lessons Learned From Animal Models. Front Immunol. 2019;10:2569.

2. Nicolai L, Leunig A, Brambs S, Kaiser R, Weinberger T, Weigand M, et al. Immunothrombotic Dysregulation in COVID-19 Pneumonia Is Associated With Respiratory Failure and Coagulopathy. Circulation. 2020;142(12):1176–89.

3. Khismatullin RR, Ponomareva AA, Nagaswami C, Ivaeva RA, Montone KT, Weisel JW, et al. Pathology of lung-specific thrombosis and inflammation in COVID-19. J Thromb Haemost. 2021;19(12):3062–72.

4. McFadyen JD, Stevens H, and Peter K. The Emerging Threat of (Micro)Thrombosis in COVID-19 and Its Therapeutic Implications. Circ Res. 2020;127(4):571–87.

5. Taquet M, Skorniewska Z, Hampshire A, Chalmers JD, Ho LP, Horsley A, et al. Acute blood biomarker profiles predict cognitive deficits 6 and 12 months after COVID-19 hospitalization. Nat Med. 2023;29(10):2498–508.

6. Brown DE, Libby SJ, Moreland SM, McCoy MW, Brabb T, Stepanek A, et al. Salmonella enterica causes more severe inflammatory disease in C57/BL6 Nramp1G169 mice than Sv129S6 mice. Vet Pathol. 2013;50(5):867–76.

7. Li J, Claudi B, Fanous J, Chicherova N, Cianfanelli FR, Campbell RAA, et al. Tissue compartmentalization enables Salmonella persistence during chemotherapy. Proc Natl Acad Sci U S A. 2021;118(51).

8. Jones AM, Mann J, and Braziel R. Human plague in New Mexico: report of three autopsied cases. J Forensic Sci. 1979;24(1):26–38.

9. Chang R, Wu DK, Wei JC, Yip HT, Hung YM, and Hung CH. The Risk of Subsequent Deep Vein Thrombosis and Pulmonary Embolism in Patients with Nontyphoidal Salmonellosis: A Nationwide Cohort Study. Int J Environ Res Public Health. 2020;17(10).

10. de Jong HK, Parry CM, van der Vaart TW, Kager LM, van den Ende SJ, Maude RR, et al. Activation of coagulation and endothelium with concurrent impairment of anticoagulant mechanisms in patients with typhoid fever. J Infect. 2018;77(1):60–7.

11. Wain J, Diep TS, Ho VA, Walsh AM, Nguyen TT, Parry CM, et al. Quantitation of bacteria in blood of typhoid fever patients and relationship between counts and clinical features, transmissibility, and antibiotic resistance. J Clin Microbiol. 1998;36(6):1683–7.

12. Hitchcock JR, Cook CN, Bobat S, Ross EA, Flores-Langarica A, Lowe KL, et al. Inflammation drives thrombosis after Salmonella infection via CLEC-2 on platelets. J Clin Invest. 2015;125(12):4429–46.

13. Beristain-Covarrubias N, Perez-Toledo M, Flores-Langarica A, Zuidscherwoude M, Hitchcock JR, Channell WM, et al. Salmonella-induced thrombi in mice develop asynchronously in the spleen and liver and are not effective bacterial traps. Blood. 2019;133(6):600–4.

14. Bevilacqua MP, Pober JS, Mendrick DL, Cotran RS, and Gimbrone MA, Jr. Identification of an inducible endothelial-leukocyte adhesion molecule. Proc Natl Acad Sci U S A. 1987;84(24):9238–42.

15. Nawroth PP, and Stern DM. Modulation of endothelial cell hemostatic properties by tumor necrosis factor. J Exp Med. 1986;163(3):740–5.

16. Herbert JM, Savi P, Laplace MC, and Lale A. IL-4 inhibits LPS-, IL-1 beta- and TNF alpha-induced expression of tissue factor in endothelial cells and monocytes. FEBS Lett. 1992;310(1):31–3.

17. Ritis K, Doumas M, Mastellos D, Micheli A, Giaglis S, Magotti P, et al. A novel C5a receptor-tissue factor cross-talk in neutrophils links innate immunity to coagulation pathways. J Immunol. 2006;177(7):4794–802.

18. Kim SJ, Carestia A, McDonald B, Zucoloto AZ, Grosjean H, Davis RP, et al. Platelet-Mediated NET Release Amplifies Coagulopathy and Drives Lung Pathology During Severe Influenza Infection. Front Immunol. 2021;12:772859.

19. Parry GC, Erlich JH, Carmeliet P, Luther T, and Mackman N. Low levels of tissue factor are compatible with development and hemostasis in mice. J Clin Invest. 1998;101(3):560–9.

20. Falati S, Liu Q, Gross P, Merrill-Skoloff G, Chou J, Vandendries E, et al. Accumulation of tissue factor into developing thrombi in vivo is dependent upon microparticle P-selectin glycoprotein ligand 1 and platelet P-selectin. J Exp Med. 2003;197(11):1585–98.

21. Wong DJ, Park DD, Park SS, Haller CA, Chen J, Dai E, et al. A PSGL-1 glycomimetic reduces thrombus burden without affecting hemostasis. Blood. 2021;138(13):1182–93.

22. Culemann S, Knab K, Euler M, Wegner A, Garibagaoglu H, Ackermann J, et al. Stunning of neutrophils accounts for the anti-inflammatory effects of clodronate liposomes. J Exp Med. 2023;220(6).

23. Payne H, Ponomaryov T, Watson SP, and Brill A. Mice with a deficiency in CLEC-2 are protected against deep vein thrombosis. Blood. 2017;129(14):2013–20.

24. Kirby AC, Yrlid U, and Wick MJ. The innate immune response differs in primary and secondary Salmonella infection. J Immunol. 2002;169(8):4450–9.

25. Mastroeni P, Villarreal-Ramos B, and Hormaeche CE. Effect of late administration of anti-TNF alpha antibodies on a Salmonella infection in the mouse model. Microb Pathog. 1993;14(6):473–80.

26. Siegmund D, and Wajant H. TNF and TNF receptors as therapeutic targets for rheumatic diseases and beyond. Nat Rev Rheumatol. 2023;19(9):576–91.

27. Qiu P, Cui X, Sun J, Welsh J, Natanson C, and Eichacker PQ. Antitumor necrosis factor therapy is associated with improved survival in clinical sepsis trials: a meta-analysis. Crit Care Med. 2013;41(10):2419–29.

28. Xie X, Li F, Chen JW, and Wang J. Risk of tuberculosis infection in anti-TNF-alpha biological therapy: from bench to bedside. J Microbiol Immunol Infect. 2014;47(4):268–74.

29. Asaduzzaman M, Rahman M, Jeppsson B, and Thorlacius H. P-selectin glycoprotein-ligand-1 regulates pulmonary recruitment of neutrophils in a platelet-independent manner in abdominal sepsis. Br J Pharmacol. 2009;156(2):307–15.

30. Wang XL, Deng HF, Tan CY, Xiao ZH, Liu MD, Liu K, et al. The role of PSGL-1 in pathogenesis of systemic inflammatory response and coagulopathy in endotoxemic mice. Thromb Res. 2019;182:56–63.

31. Opal SM, Sypek JP, Keith JC, Jr., Schaub RG, Palardy JE, and Parejo NA. Evaluation of the safety of recombinant P-selectin glycoprotein ligand-immunoglobulin G fusion protein in experimental models of localized and systemic infection. Shock. 2001;15(4):285–90.

32. Kum WW, Lee S, Grassl GA, Bidshahri R, Hsu K, Ziltener HJ, et al. Lack of functional P-selectin ligand exacerbates Salmonella serovar typhimurium infection. J Immunol. 2009;182(10):6550–61.

33. Dalton DK, Pitts-Meek S, Keshav S, Figari IS, Bradley A, and Stewart TA. Multiple defects of immune cell function in mice with disrupted interferon-gamma genes. Science. 1993;259(5102):1739-42.

34. Finney BA, Schweighoffer E, Navarro-Nunez L, Benezech C, Barone F, Hughes CE, et al. CLEC-2 and Syk in the megakaryocytic/platelet lineage are essential for development. Blood. 2012;119(7):1747–56.

35. Peschon JJ, Torrance DS, Stocking KL, Glaccum MB, Otten C, Willis CR, et al. TNF receptor-deficient mice reveal divergent roles for p55 and p75 in several models of inflammation. J Immunol. 1998;160(2):943–52.

36. McDonald B, Davis RP, Kim SJ, Tse M, Esmon CT, Kolaczkowska E, et al. Platelets and neutrophil extracellular traps collaborate to promote intravascular coagulation during sepsis in mice. Blood. 2017;129(10):1357–67.

