## Supplemental material for "Discrete and conserved inflammatory signatures drive thrombosis in different organs after *Salmonella* infection"

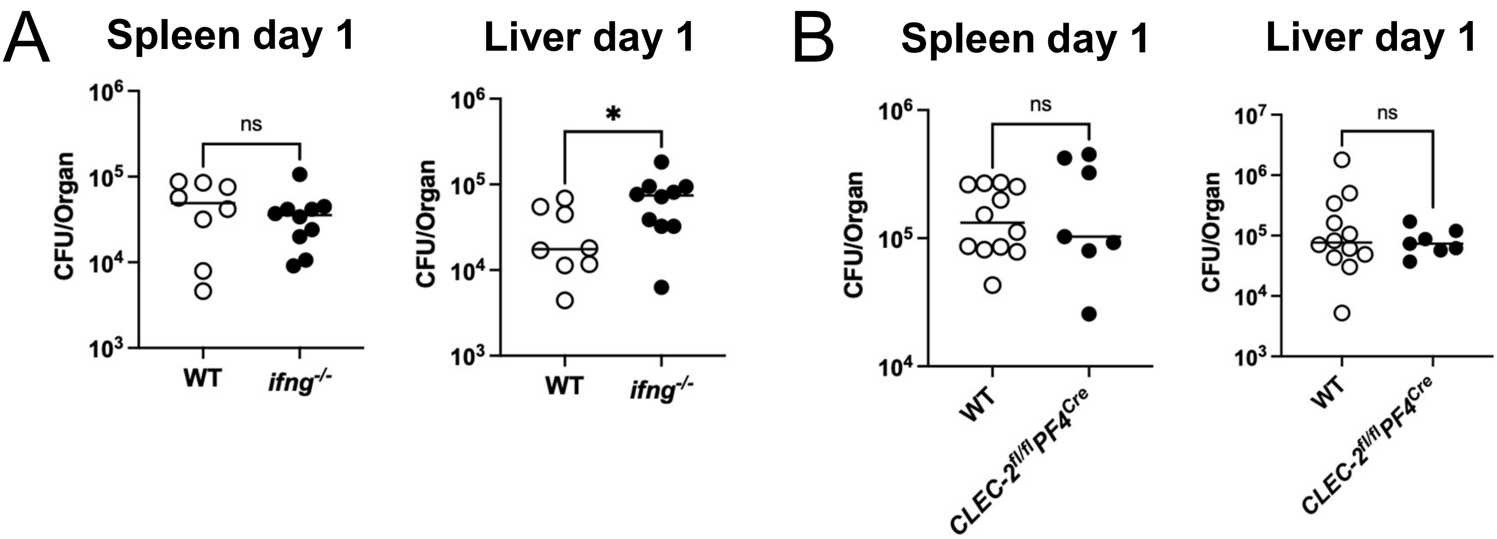


**Supplementary figure 1**. Colony Forming Units (CFU) per spleen and liver after 24 h of infection with STm. Mice were infected with $5x10^5$ CFU STm SL3261. (A) WT or IFN-γ-deficient mice (B) WT or CLEC-$2^{fl/f}$lPF4$^{Cre}$ mice. Each dot represents an individual mouse. Horizontal lines depict the median. Mann-Whitney test was applied. *P<0.05, ns=non-significant.

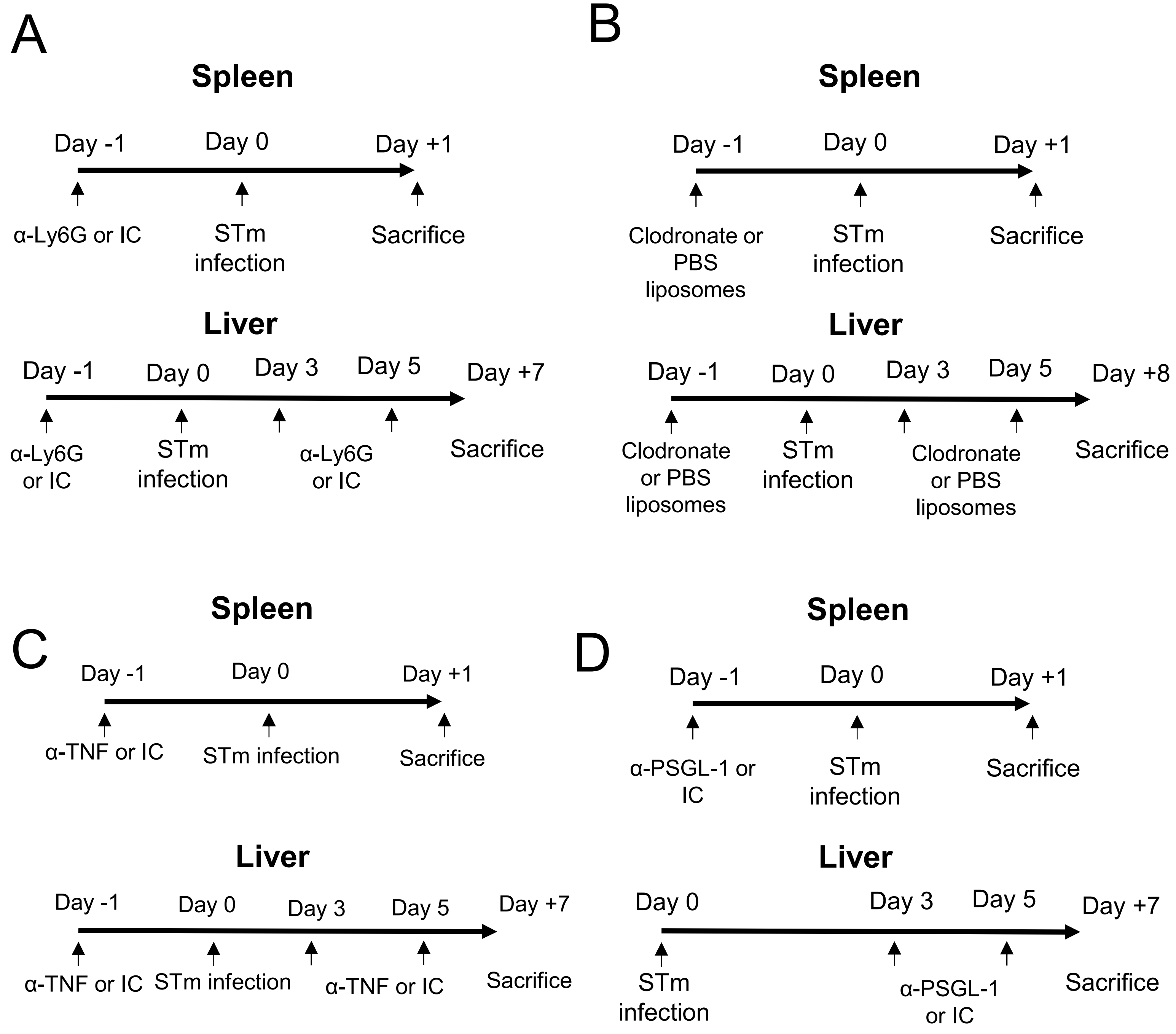


**Supplementary figure 2**. Experimental approach for the depletion of neutrophils (A), monocytic lineage cells (B), blocking TNF (C) or blocking PSGL-1 (D). Briefly, 6-8 weeks old C57Bl/6 mice were infected with $5x10^5$ CFU i.p. with STm SL3261. Antibodies or controls were administered via i.p. in the time points indicated.

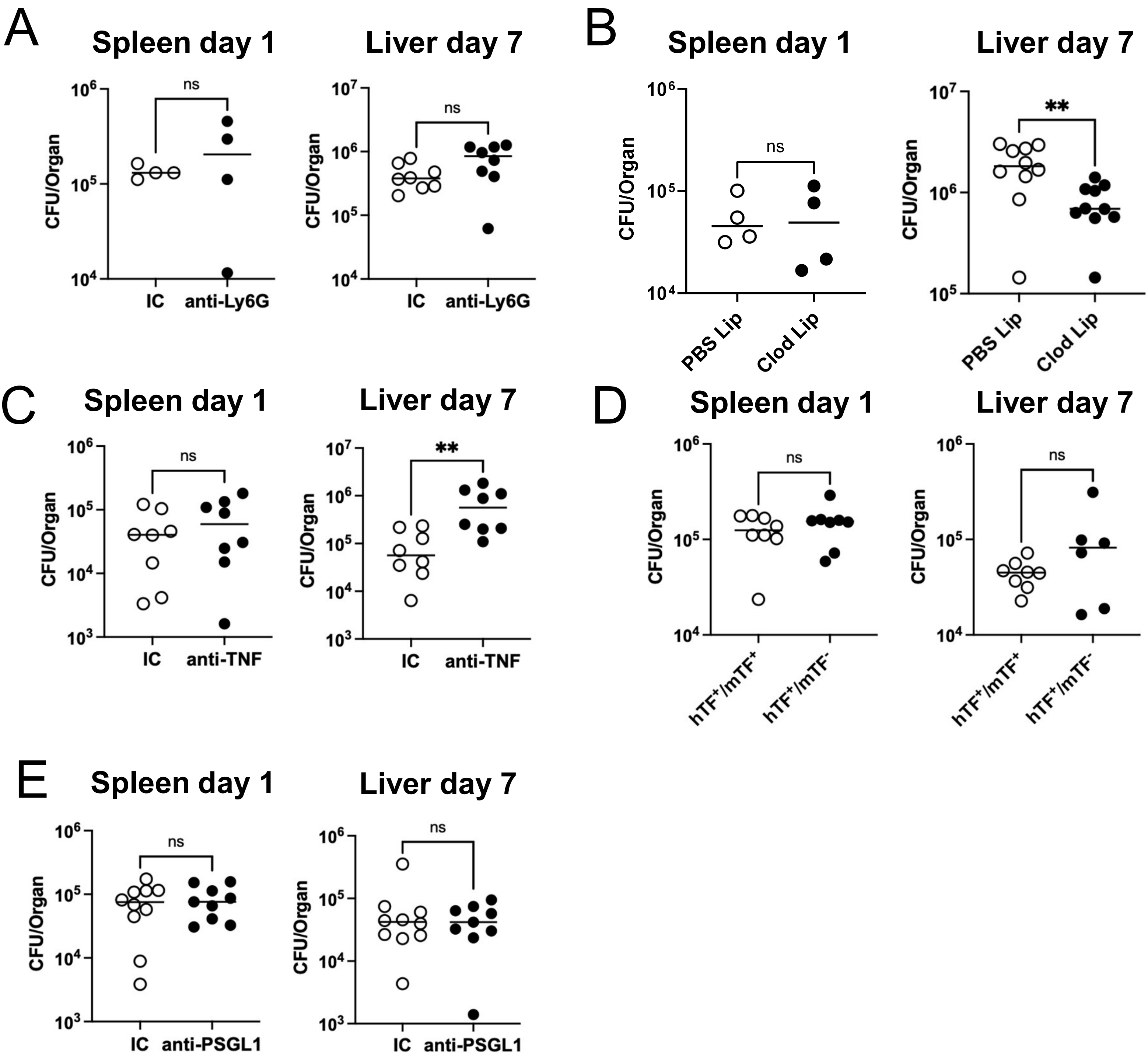


**Supplementary figure 3**. Colony Forming Units (CFU) per spleen after 24 h, or liver after 7 days of infection. Mice were infected with $5x10^5$ CFU STm SL3261. (A) isotype control or anti-Ly6G treated mice, (B) PBS liposomes or clodronate liposomes treated mice, (C) Isotype control or anti-TNF treated mice, (D) isotype control or anti-PSGL-1 mice and (E) TF-sufficient controls ($hTF^+/mTF^+$) or low TF mice ($hTF^+/mTF^-$). Each dot represents an individual mouse. Horizontal lines depict the median. Mann-Whitney test was applied. ns=non-significant.

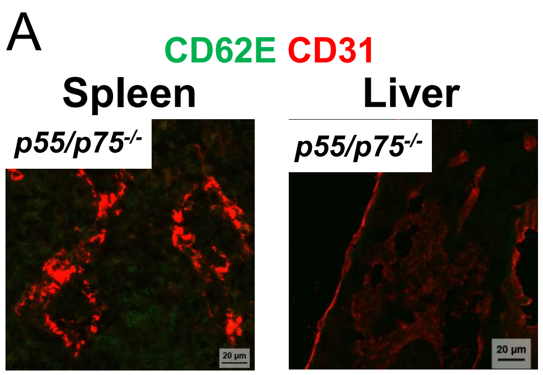


**Supplementary figure 4**. (A) Representative images of spleen and liver sections from mice deficient in the TNF receptors p55 and p75, infected with STm as previously described. Green is CD62E and red is CD31. n=4

### Supplementary table I. Reagents used in this study

| Target | Format | Manufacturer | Host | Clone | Cat No. |
|---|---|---|---|---|---|
| ***Primaries*** | | | | | |
| CD41 | Purified | eBioscience | Rat | eBioMWReg30 | 14-0411-85 |
| Fibrin | Purified | Accurate chemical & Scientific corporation | Goat | Polyclonal | |
| Ly6G | Purified | BD Pharmingen | Rat | 1A8 | 551459 |
| Ly6G | Biotin | Biolegend | Rat | 1A8 | 127604 |
| F480 | plexa Fluor 488 | Invitrogen | Rat | BM8 | 14-4801-82 |
| F480 | blexa Fluor 488 | Biolegend | Rat | BM8 | 123106 |
| Ly6C | Biotin | Biolegend | Rat | HK1.4 | 128004 |
| TNF | Purified | Abcam | Rabbit | Polyclonal | ab9739 |
| CD31 | plexa Fluor 488 | Invitrogen | Rat | 390 | 14-0311-82 |
| CD31 | Biotin | Invitrogen | Rat | 390 | 13-0311-83 |
| CD62E | E | BD Pharmingen | Rat | 10E9.6 | 550290 |
| aSMA | Cy3 | Sigma | Mouse | 1A4 | C6198 |
| TF | Purified | R&D systems | Goat | polyclonal | AF3178 |
| CD11b | blexa Fluor 488 | eBioscience | Rat | M1/70 | 17-0112-81 |
| PSGL-1 | Purified | BD Pharmingen | Rat | 2PH1 | 564310 |
| ***Secondaries*** | | | | | |
| anti-rat | Biotin | Dako | Rabbit | Polyclonal | E0468 |
| Anti-sheep | HRP | Jackson Immunoresearch | Donkey | Polyclonal | 713035147 |
| anti-rabbit | Alexa Fluor 488 | Jackson Immunoresearch | Donkey | Polyclonal | 711-545-152 |
| anti-AF488 | Alexa Fluor 488 | Invitrogen | Rabbit | Polyclonal | 710369 |
| anti-rat | Alexa Fluor 647 | Jackson Immunoresearch | Donkey | Polyclonal | 712-605-153 |
| Anti-rat | Cy3 | Jackson Immunoresearch | Donkey | Polyclonal | 712-165-153 |
| Anti-sheep | Alexa Fluor 488 | Jackson Immunoresearch | Donkey | Polyclonal | 713-545-147 |
| Vectastain® ABC-Alkaline Phosphatase | Alkaline Phosphatase | Vector Laboratories | | | AK-5000 |
| AF555 Streptavidin | Alexa Fluor 555 | Invitrogen | | | S32355 |
| ***IVM*** | | | | | |
| CD49b | PE | Biolegend | Arm Hms | HMα2 | 103506 |
| Ly6G | Brilliant Violent 421 | Biolegend | Rat | 1A8 | 127628 |
| F480 | Alexa Fluor 647 | Biolegend | Rat | BM8 | 123122 |
| Thrombin Activity Assay | | Anaspec | | | AS-72129 |
| ***Flow Cytometry*** | | | | | |
| Zombie Aqua Viability dye | Brilliant Violent 510 | Biolegend | | | 423101 |
| CD45 | PE-Cy7 | | Rat | 30-F11 | 103114 |
| CD11b | Alexa Fluor 700 | BD Pharmingen | Rat | M1/70 | 557960 |
| Ly6G | PE CF594 | BD Horizon | Rat | 1A8 | 562700 |
| Ly6G | APC | BD Pharmingen | Rat | 1A8 | |
| Ly6C | PerCP Cy5.5 | eBioscience | Rat | HK1.4 | 25-5932-82 |
| F480 | Brilliant Violent 421 | Biolegend | Rat | BM8 | 123137 |
| CD3 | Alexa Fluor 488 | eBioscience | Rat | 145-2C11 | 53-0031-82 |
| B220 | FITC | BD Pharmingen | Rat | RA3-6B2 | 553087 |
| NK1.1 | FITC | BD Pharmingen | Rat | PK136 | 553164 |
| TNF | PE Dazzle 594 | Biolegend | Rat | MP6-XT22 | 506346 |
| Cytofix/Cytoperm Fixation/permeabilization kit | | | | | 554714 |
| ***Blocking/depleting reagents*** | | | | | |
| Invivo Mab anti-mouse Ly6G | Purified | BioXcell | Rat | | BE0075-1 |
| Clodronate Liposomes | | Liposoma | | | C-005 |
| TNF | Purified | BioXcell | Rat | XT3.11 | BE0058 |
| Invivo Mab anti-mouse PSGL-1 (CD162) | Purified | BioXcell | Rat | 4RA10 | |
| Rat IgG | Purified | Sigma | Rat | Polyclonal | I4131 |
